# Long-term performance of disease-resistant grapevine varieties: insights from an 8-year field monitoring across French vineyards

**DOI:** 10.64898/2026.04.28.721351

**Authors:** Rémi Pelissier, Laura Marolleau, Isabelle D. Mazet, François Delmotte, Laurent Delière, Anne-Sophie Miclot, Frédéric Fabre

**Affiliations:** INRAE, Bordeaux Sciences Agro, SAVE, ISVV, F-33140 Villenave d’Ornon, France

**Keywords:** Sustainable agriculture, plant genetic resistance, resistance durability, grapevine, pathogen field monitoring

## Abstract

Breeding disease-resistant varieties (DRV) is a central strategy for reducing reliance on phytosanitary products. However, the successful deployment and long-term durability of these cultivars rely on acquiring field data across diverse production conditions, a step that remains frequently neglected, especially in perennial crops. Since 2018, the OSCAR observatory, a network of vineyard plots planted in France with varieties resistant to downy and powdery mildew, the two major pathogens of grapevine, has aimed to close this gap for viticulture. The observatory comprises over 199 commercial plots, covering 127 hectares across diverse agroclimatic conditions, all managed by winegrowers under their own production practices. The observatory currently monitors 30 disease-resistant grapevine varieties, tracking both their agronomic performance and the dynamics of key pathogens. Since 2018, while phytosanitary treatments have been reduced by an average of 79% compared to conventional plots, the incidence of downy and powdery mildew, remain low, even in years highly conducive to these diseases. However, the long-term survey also highlights the decline in efficacy of some resistances to downy mildew and the emergence of black rot, a disease effectively controlled by conventional phytosanitary programs. Beyond acting as a rapid warning system for resistance breakdown, the observatory promotes sustainable disease management in viticulture. It provides valuable insights to winegrowers on effective DRV management. It also delivers actionable feedback to breeders to guide more durable DRV breeding strategies.

**Highlights:**

- OSCAR observatory monitors 199 plots of grapevine disease resistant varieties (DRV)
- Grapevine DRV cuts fungicide uses by 79% while maintaining good disease control
- Some resistances efficacy declines against downy mildew, but not powdery mildew
- Black rot, a disease usually controlled by fungicide, is rising in OSCAR plots OSCAR provides useful feedback to breeders and winegrowers on DRV management

## Introduction

Plant pathogens are responsible for significant crop loss and represent an increasing threat to global food security (Savary et al., 2019; Singh et al., 2023; Teng et al., 1984). At the same time, societal and regulatory pressures are pushing to reduce the use of chemical plant protection products (Young et al., 2022). The adoption of disease-resistant crop varieties is a key lever to address this demand. However, breeding resistant varieties remains a long and costly process, and their effectiveness is often limited in field conditions as pathogens may often adapt rapidly to newly introduced resistances (Parlevliet, 2002; Peressotti et al., 2010; Rimbaud et al., 2021). Accordingly, the deployment of resistant varieties must be supported by field monitoring to provide technical recommendations facilitating their integration into cropping systems in the short term and to promote resistance durability in the longer term (Soubeyrand et al., 2024).

The epidemiological control achieved in the field by resistant cultivars largely depends on the qualitative or quantitative phenotypic expression of resistance genes (Niks et al., 2015). Complete (i.e. qualitative) resistance, typically conferred by major resistance genes, fully suppress the disease. Often mediated by gene-for-gene interactions, qualitative resistances remain effective as long as the corresponding avirulence allele is present in pathogen populations. In case of resistance breakdown, the epidemiological control is usually lost abruptly (Mundt, 2014). In contrast, partial (i.e. quantitative) resistance reduces disease severity by limiting pathogen development on the host. While long considered to be polygenic (French et al., 2016), partial resistance can also result from a single major gene with an incomplete effect on disease reduction (Cowger and Brown, 2019; Niks et al., 2015). Pathogen adaptation to partial, polygenic resistance is commonly referred to as erosion, while adaptation to monogenic, quantitative resistance is called breakdown, consistent with the terminology used for qualitative resistance (Hammond-Kosack and Jones, 1997; Jones and Dangl, 2006).

The durability of resistant cultivars in the field depends on a complex interplay of factors, including the design choices made by breeders (e.g., selection of R genes), their spatial and temporal deployment, agricultural management practices, pathogen biology, and environmental conditions (Rimbaud et al., 2021; Zaffaroni et al., 2023). For example, resistance based on a single gene (monogenic resistance) is generally less durable than pyramided resistance, since the simultaneous adaptation of a pathogen to several resistance factors is more complex (Gandon et al., 2024; Mundt, 2018; Pilet-Nayel et al., 2017). Pathogens combining sexual and asexual reproduction and capable of adapting to a wide range of environmental conditions exhibit a higher evolutionary potential to overcome host resistance (McDonald and Linde, 2002). More generally, the biology of the host-pathogen system strongly interacts with the deployment strategies of resistant cultivars in space and time to determine resistance durability (Burdon et al., 2014; Zaffaroni et al., 2023). In annual cropping systems, agronomic practices such as crop/varieties rotation, delayed fall planting, varietal mixture or tillage allows to take advantage of counter-selection mechanisms to extend the lifespan of a resistance once it has been overcome (Burdon et al., 2014; Rimbaud et al., 2018). In contrast, perennial crops such as fruit trees or grapevines, offer far lever to manage resistance breakdown (REX Consortium et al., 2016), making the deployment and management of resistant cultivars a critical component of their long-term durability (Cox et al., 2005).

Mathematical modelling can be used to compare the epidemiological, evolutionary, and economic performance of deployment strategies for perennial plant in order to provide mechanistic insight into the consequences of different deployment strategies, and allow decision-makers to understand their relative merits (Rimbaud et al., 2021; Zaffaroni et al., 2025). However, most models developed to date remain primarily conceptual and are not designed to generate quantitative predictions under real-world conditions. Bridging this gap requires a close, interactive dialogue between modelling and empirical evidence, in which field-based observational and experimental data are used to parameterize, validate, and refine models, while model outputs in turn inform data collection and monitoring priorities. In the short to medium term, observational approaches remain the only means to document the effective potential of disease-resistant varieties under real production situations (i.e. the set of physical, biological, and socioeconomic factors that jointly determine agricultural production; Savary et al., 2019).

In this context, field monitoring networks spanning diverse production situations are essential, not only to document resistance performance, but also to anchor modelling frameworks in the complexity of real agroecosystems. As demonstrated by some rare initiatives such as the Vinquest project, which has monitored the impact of apple resistance genes against apple scab for more than ten years across Europe, the durability of varietal resistance varies considerably between resistance genes and production situations (Patocchi et al., 2020). By facilitating the early detection of resistance breakdowns, this project has enabled to prioritize in breeding programs the introgression of resistance genes that remain effective under field conditions. Lack of monitoring delays the detection of resistance breakdown until the virulent pathotype is already widespread in pathogen population. Such a detrimental situation has for example occurred in poplar rust where resistant poplar carrying the *RMlp7* gene were still planted more than nine years after the first breakdowns, leading to the development of virulent pathogen populations (Louet et al., 2023). Similarly, resistance management against coffee leaf rust, caused by *Hemileia vastatrix*, is increasingly challenging (Cabral et al., 2009) due to the widespread breakdowns and the emergence of strains combining virulence factors against most resistance genes (Alves et al., 2024). As a result, there is a renewed reliance on chemical control strategies rather than on varietal resistance (Sera et al., 2022).

An original initiative for the in-field assessment of resistance durability is the OSCAR observatory, dedicated to monitoring grapevine Disease Resistant Varieties (DRVs) under real world viticultural practices in France. The OSCAR observatory was launched in 2017 alongside the registration of the first resistant varieties in the French national catalog. It now includes over 30 distinct resistant cultivars, distributed across nearly all French wine-growing regions (Guimier et al., 2019; Merdinoglu et al., 2018). Grapevine DRVs carry different genetic resistance combination, either monogenic or pyramided, involving *Rpv* (Resistance to *Plasmopara viticola*), *Run*, and *Ren* (Resistance to *Uncinula/Erysiphe necator)* factors. Currently, only *Rpv* factors which confer partial resistance against downy mildew are used in commercial grapevine DRV while those use against powdery mildew confer either total (*Run*) or partial (*Ren*) resistance (Possamai and Wiedemann-Merdinoglu, 2022). These diseases are the two most destructive diseases of grapevine, making viticulture one of the most fungicide- intensive cropping systems (Gessler et al., 2011; Koledenkova et al., 2022; Nefti et al., 2024). Each year many variables are collected in the OSCAR observatory, including data on major and secondary grapevine diseases, technical practices, and the agronomic performance of each variety. Here, we analyzed the dataset gathered during eight years of monitoring to address two research questions. First, how does grapevine DRVs cultivation impact phytosanitary treatments and the epidemiology of grapevine diseases? Second, how effective and durable are the resistance deployed against powdery and downy mildews?

## Material and methods

### Data collection

The OSCAR observatory monitors vineyard plots planted with grapevine disease resistant varieties (DRVs) to downy mildew (DM; caused by *Plasmopara viticola*) and powdery mildew (PM; caused by *Erysiphe necator*). These varieties originate from European breeding programs conducted in Italy, Germany, and Switzerland (Salotti et al., 2022) and in France by INRAE (Merdinoglu et al., 2018). All data used in this study were retrieved from the OSCAR observatory database (see Results for a description of the network, or Guimier et al., 2019).

The plots are distributed across a wide range of agroclimatic conditions and managed by winegrowers according to their own production objectives. The plots included in the network must cover at least 0.2 ha (approximately 700–1500 vines), be planted with a single grapevine DRV, and follow vineyard management practices consistent with those applied to the rest of the estate. Each plot is characterized by its geographical location, plot surface area, cultivar, year of planting, planting density, and rootstock. Every year, agronomic and plant health variables are recorded. They include the management practices (notably phytosanitary treatments), the impact of ten major grapevine diseases and the disease pressure observed in surrounding plots cultivated with susceptible grapevine varieties. Every year, the following agronomic data are collected:

- Plant protection practices: for each treatment, the date, target disease, product name and active ingredients, applied dose, and decision rules are recorded. These data are used to calculate the Treatment Frequency Index (TFI). The fungicide TFI was calculated as the sum, for all fungicide applications (including biocontrol products), of the applied dose divided by the recommended registered dose (Pingault et al., 2009). Fungicide TFIs are separated from insecticide and herbicide treatments for subsequent analyses.
- Assessment of local disease pressure: disease pressure for downy mildew, powdery mildew, and black rot is evaluated on susceptible *Vitis vinifera* cultivars growing close to the monitored plot. Disease pressure is scored as absence of disease (0), low (1), intermediate (2), or high (3).
- Assessment of disease scores for ten major grapevine diseases: standardized protocols are used to record data on the impact of diseases both on leaves, clusters or stem. Disease assessments are performed between the phenological stage of veraison until the harvest stage (Coombe, 1995), once all phytosanitary treatments have been completed. The diseases and pests surveyed are DM and PM (the target of resistance genes), Black Rot (BR) caused by *Guignardia bidwellii*, Anthracnosis caused by *Elsinoë ampelina*, Phomopsis cane caused by *Diaporthe ampelina*, Gray mold caused by *Botrytis cinerea*, Erineum caused by the mite *Colomerus vitis*, Phylloxera caused by *Daktulosphaira vitifoliae*, Sour rot (caused by a complex of yeasts and bacteria, often associated with *Drosophila spp*.*) and* Leafhopper burn caused by leafhoppers *(e*.*g. Empoasca vitis)*. For each pest and disease, incidence and severity are assessed at the plot scale on leaves and clusters (or stem for Phomopsis cane), using a semi-quantitative scale ranging from 0 to 5 (**Table 1**).

**Table 1.**
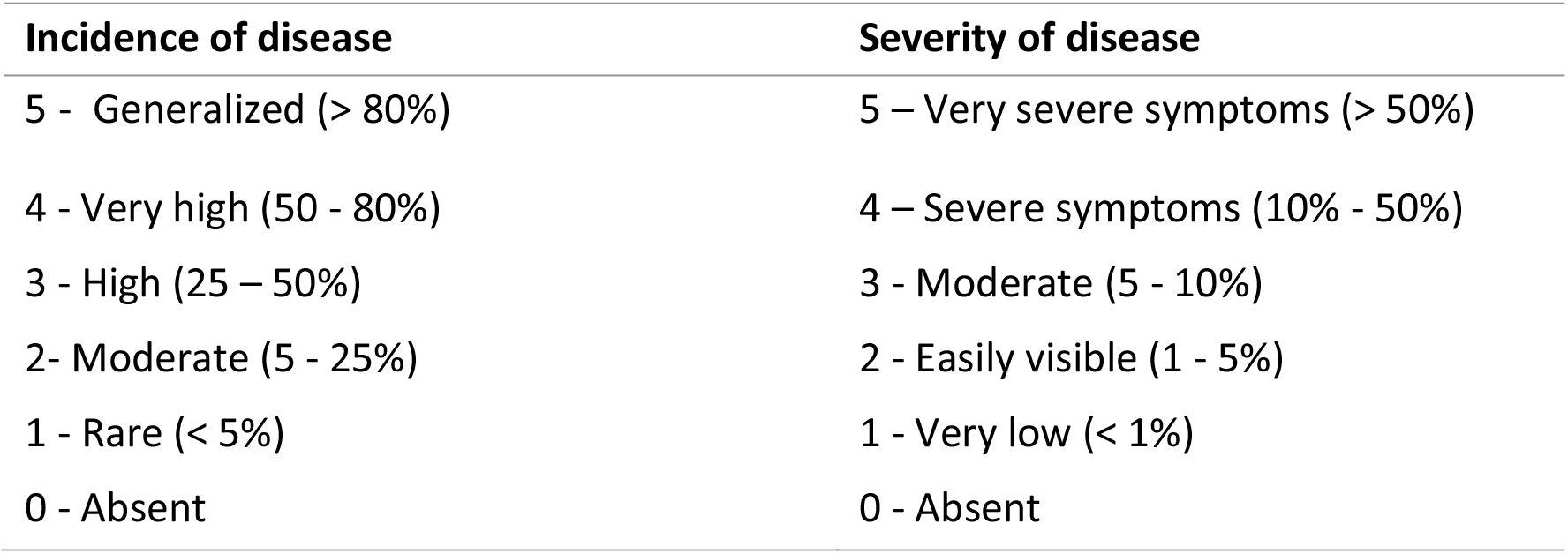
Rating scales used for pest and disease assessments on OSCAR plots. Four variables are evaluated at the plot scale: incidence on leaves and clusters (frequency of vines with at least one symptomatic leaf and frequency of vines with at least one symptomatic cluster) and severity on leaves and clusters. Severity evaluation includes healthy and diseased organs (overall percentage of the surface of leaves or clusters with symptoms). For example, if 50% of the leaves from a plot are affected by downy mildew (incidence) with an average severity of 25% for those leaves, the severity of the plot will be 50%*25% = 12,5%.

For each variable, five classes have been defined:

- Incidence on leaves and clusters (plot scale): 5: Generalized (>80%), 4: Very high (50- 80%), 3: High (25-50%), 2: Moderate (5-25%), 1: Rare (<5%), 0: Absent.
- Severity (plot scale) 5: Very high (>50%), 4: High (10-50%), 3: Moderate (5-10%), 2: Easily visible (1-5%), 1: Very low (<1%) 0: Absent.

### Statistical analysis and representation

The annual mean TFI in the OSCAR observatory was compared to the national TFI reference, calculated from the Agreste 2019 database, which reports average TFIs for susceptible grapevine varieties in France. This national value was used as a reference to perform a two-sided Student’s t-test, testing whether the annual mean fungicide TFI in OSCAR observatory was significantly different than the national reference. The distribution of incidence (frequency of diseased plants) and severity (intensity of symptoms) classes of each disease across plots were visualized for year and organ as stacked histograms (**Figure 2** for DM and PM, and **Figure 5** for BR). For all other grapevine diseases, only the maximum incidence value recorded for either leaves or clusters was represented (**Figure 4**).

### Statistical analysis

Statistical analyses were conducted to identify the factors influencing foliar disease severity. Severity scores were selected because they integrate both the proportion of affected organs and symptom severity; leaf assessments were preferred because they are more prevalent than cluster assessments due to the young age of the plots in the OSCAR observatory. For each disease (DM, PM and BR) severity scores were transformed into a binary response variable at the plot level. Scores of 0 and 1 (<1% severity) were classified as low disease severity, whereas scores ranging from 2 to 5 (>1% diseased plants) were classified as high disease severity.

The resulting binary response variable was analyzed using a binomial generalized linear model (GLM) with a logit link function. Due to differences in the genetic resistance factors available for each disease, two distinct full model structures were fitted to evaluate the relative importance of climatic, agronomic, and epidemiological variables.

The effect of the explanatory variables on the probability of high disease severity (*i*.*e*. severity > 1%) was assessed using an analysis of deviance on the selected models. The significance of the main effects and their interaction was evaluated using a Type II ANOVA. To rank the relative importance of each explanatory variable, we computed the partial deviance explained by degrees of freedom, expressed as a percentage of the null deviance. This metric (called hereafter “Relative importance”) is analogous to a partial pseudo-R^2^ scaled by degrees of freedom. For DM and PM, differences between resistance factors were tested via pairwise comparisons using Tukey’s HSD post-hoc tests and visualized by plotting fitted probabilities extracted from the models. An identical approach was applied to assess the effect of the DRV on the probability of high BR severity.

### Model structure for downy mildew (DM) and powdery mildew (PM)

The full model included the following explanatory variables: survey year (8 levels; categorical), agro-climatic context (Oceanic vs. Mediterranean; categorical, plot localized in center and north east France were excluded due to low representation) (**Figure 1**), local disease pressure (continuous), fungicide use (total fungicide TFI applied prior to assessment; continuous), plot age (continuous), deployed resistance factor (5 levels for DM and 3 levels for PM; categorical), and the two-way interaction between plot age and resistance factor. To ensure statistical robustness, only resistance factors present in at least three plots (*Rpv1, Rpv1-Rpv3*.*1, Rpv3*.*2, Rpv10, Rpv3*.*1-Rpv12*) were retained. Due to their low resistance levels, DRVs carrying *Rpv10* and *Rpv3*.*3-Rpv10* were group together (Foria et al., 2018; Wilkerson et al., 2025). Underrepresented factors (*Rpv1–Rpv10, Rpv3*.*1*, and *Rpv12*) were excluded from the analysis.

**Figure 1.**
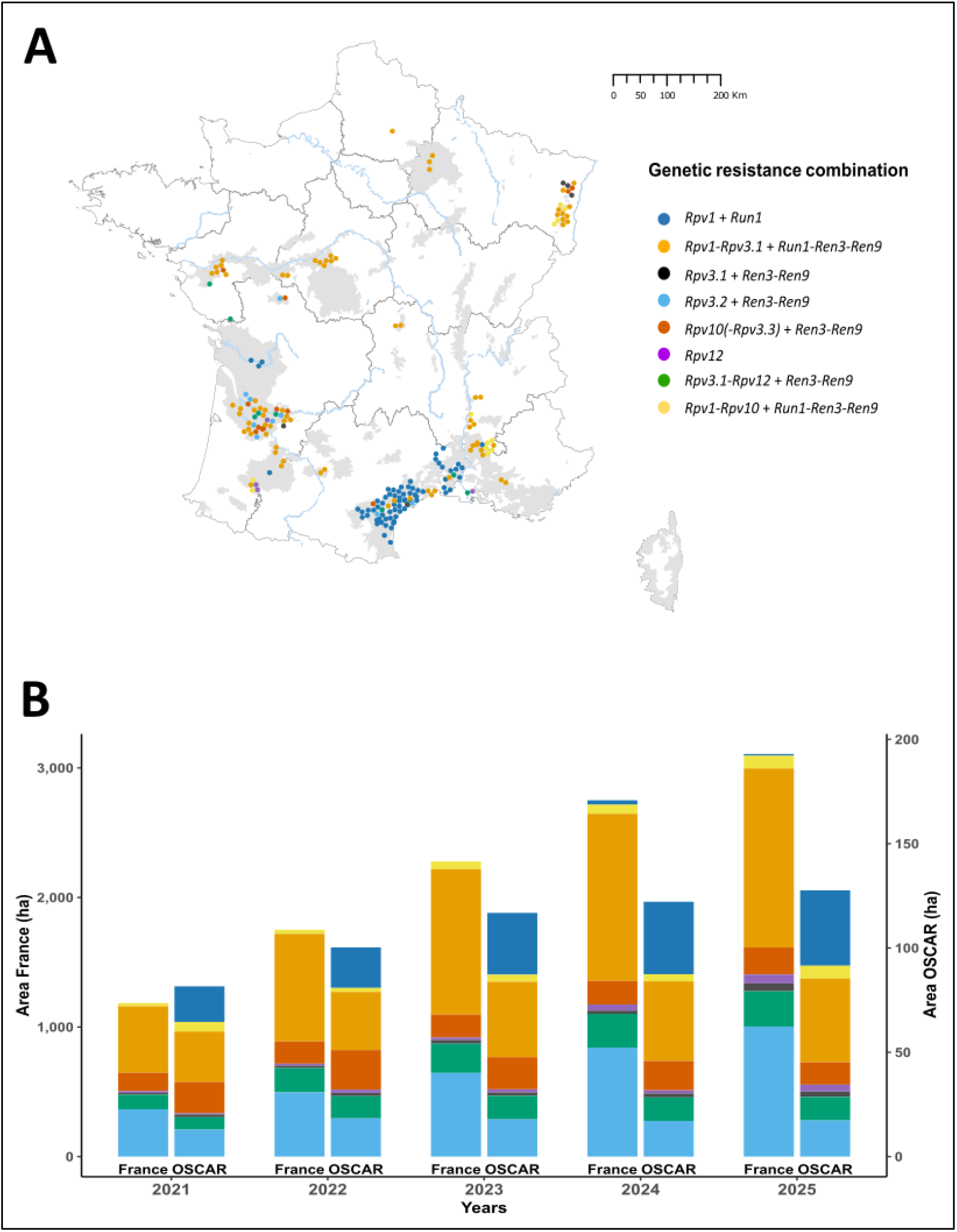
Localization, area, plots and resistance deployed in the OSCAR observatory. A) Map of the 199 plots of the OSCAR observatory in 2025. Colors indicate the genetic resistance combinations deployed against downy mildew (Rpv factors) and powdery mildew (Run or Ren factors). The number of plots used in the study and the associated varieties for each genetic resistance combination are presented in Table 2. Grey area delineates the main wine-growing region. B) Total area of grapevine genetic resistance combination planted in France (source: INAO) and monitored in the OSCAR observatory since 2021 (Rpv, Run and Ren; colors are the same as in panel A).

**Figure 2.**
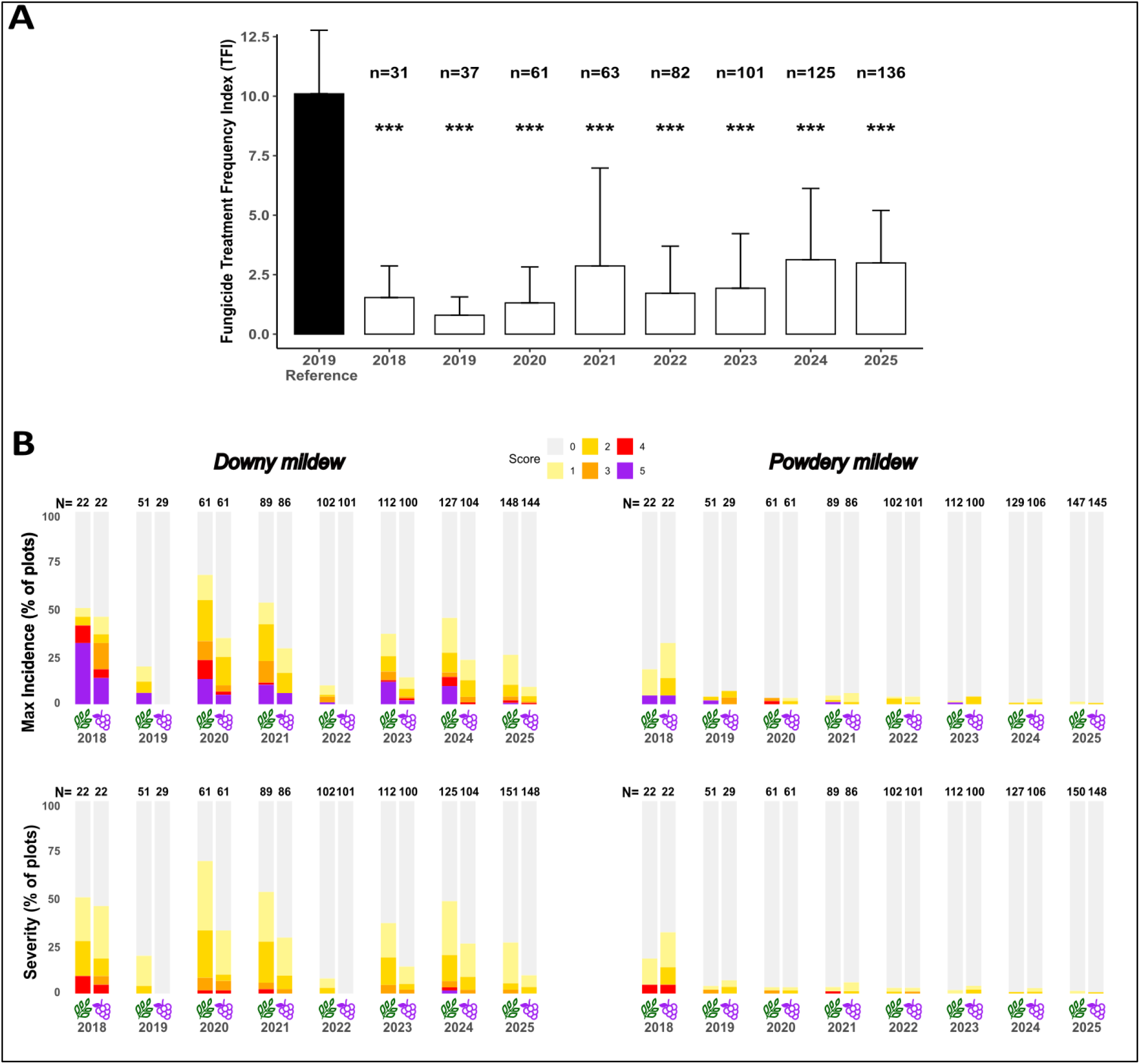
Fungicide treatment and disease notation in the OSCAR observatory. **A)** Mean of fungicide treatment frequency index (TFI) in the OSCAR observatory from 2018 to 2025 along with the national reference 2019 calculated on susceptible grapevines. All OSCAR plots are planted with grapevine resistant varieties. Stars indicate a mean of TFI significantly different from the national reference represented in black according to a two-sided Student’s t-test (*: p < 0.05; **: p < 0.01, ***: p < 0.001). **B)** Incidence (Frequency of plant with at least one symptom per plot) and Severity (Intensity of symptom) of downy mildew and powdery mildew on leaves and clusters in the plots surveyed at the veraison or harvest stages since 2018. The histograms report the percentage of the plots in each class (from 0 to 5, see methods) with sample sizes indicated for each year and organ.

To explore the interaction between plot age and resistance factors for DM, the slope of the probability of high DM severity with respect to plot age was estimated for each resistance factor. Predicted probabilities of high DM severity were computed for plot ages ranging from 1 to 13 years for each resistance factor, for each year of survey, agroclimatic condition, and setting fungicide use and disease pressure to their mean values. These predicted probabilities were visualized across three classes of plot age: ≤ 3 years, 3–6 years, and > 6 years

### Model structure for black rot (BR)

The full model included the same set of explanatory variables with three modifications. First, as no resistance factors against black rot are known in the monitored varieties, the identity of the DRV (15 levels; categorical) was used instead of the resistance factor. Only variety present in at least two different plots were kept for the analysis. Second, a binary variable describing the infectious history of the plot regarding BR (previously infected at least once vs. never infected; categorical) was included. Third, the interaction between plot age and DRV was omitted.

To investigate the interaction between plot age and infectious history for BR, predicted probabilities of BR presence were computed for plot ages ranging from 1 to 13 years, for both never infected and previously infected plots, for each year of survey, agroclimatic condition, and setting fungicide use and disease pressure to its mean value. Prediction uncertainty was quantified using a non-parametric bootstrap approach. Three hundred bootstrap samples were generated by resampling the plots with replacement; the model was refitted on each sample, and marginal predictions were recalculated. The 95% confidence intervals were subsequently derived from the 2.5th and 97.5th percentiles of the bootstrap distributions.

### Software used

Statistical analyses were performed with R software version version 2024.12.0; R 206 Core Team, 2024. GLM were fitted with the lme4 package (Bates et al., 2015). Statistical differences between levels of significant categorical explanatory variables were tested with pairwise comparisons using Tukey’s HSD post hoc tests implemented with the function glht() from the package multcomp (Bretz et al., 2008). Trend estimations for the interactions between plot age and resistance factor were performed with the function emtrends() from the package emmeans (Lenth, 2017). All plots were generated using the package ggplot2. Bootstrap samples (B = 300) were generated using a custom function, based on plot-level resampling with replacement and refitting of the binomial generalized linear model for each replicate. The data and analysis scripts used in this study are available from the authors upon request.

## Results

### OSCAR observatory

The OSCAR observatory is a network of vineyard plots planted with downy and powdery mildew (DM, PM) DRVs all across France. Established since 2017, OSCAR was launched by INRAE (French national institute of agronomy and environment) and IFV (French institute of Wine) following the first authorization of DRVs cultivation by winegrowers in France. It involves more than 30 partners institutions, including wine interprofessional organizations, Chambers of Agriculture, and the French Federation of Grapevine Nurseries.

The plots are distributed across a wide range of agroclimatic conditions and managed by winegrowers according to their own production objectives. New plots are integrated into the OSCAR observatory each year. In 2018, OSCAR comprised 32 plots across 14 locations representing 16.7 ha. By 2025, it had expanded to 199 plots, representing 127 ha, with an average plot size of 0.64 ha. The observatory currently monitors 30 mildew-resistant varieties, including 18 developed by INRAE and 12 originating from other European breeding programs (**Figure 1, Table 2**).

**Table 2.** Varieties and resistance factors monitored in the OSCAR observatory. Resistance factors for Downy and Powdery mildews are listed for each genetic resistance combination. The number of plots used in this study (available data) is indicated in bold for each combination, relative to the total number of plots recorded in the OSCAR observatory between brackets.

| Genetic combination |  |  |  |  |  |  |  |  |  |  |
| --- | --- | --- | --- | --- | --- | --- | --- | --- | --- | --- |
| DM resistance factor | PM resistance factor | Variety | 2018 | 2019 | 2020 | 2021 | 2022 | 2023 | 2024 | 2025 |
| *Rpv1* | *Run1* | Coutia, Luminan and Bouquet variety (5 cultivars) | **0** (5) | **13** (18) | **25** (29) | **29** (30) | **31** (34) | **47** (53) | **63** (68) | **67** (71) |
| *Rpv1-Rpv3.1* | *Run1-Ren3-Ren9* | Artaban, Floreal, Vidoc, Voltis | **6** (11) | **16** (29) | **15** (38) | **30** (49) | **43** (60) | **27** (65) | **30** (73) | **55** (78) |
| *Rpv3.1* | *Ren3-Ren9* | Prior, Pinotin, Johanniter | **1** (1) | **0** (1) | **0** (1) | **1** (1) | **2** (2) | **2** (2) | **3** (3) | **3** (4) |
| *Rpv3.2* | *Ren3-Ren9* | Souvignier Gris | **1** (1) | **5** (8) | **6** (9) | **7** (10) | **6** (11) | **8** (10) | **6** (8) | **7** (9) |
| *Rpv10 (+Rpv3.3)* | *Ren3-Ren9* | Cabernet Cortis, Monarch, Muscaris, Divico | **2** (6) | **4** (10) | **4** (11) | **5** (13) | **6** (15) | **7** (13) | **7** (13) | **8** (13) |
| *Rpv12* | *None* | Merlot Khorus, Cabernet Volos | **1** (1) | **1** (1) | **1** (1) | **1** (1) | **2** (2) | **2** (2) | **2** (2) | **1** (4) |
| *Rpv3.1-Rpv12* | *Ren3-Ren9* | Cabernet Jura, Sauvignac, Soreli | **5** (5) | **5** (7) | **5** (8) | **5** (7) | **4** (7) | **3** (8) | **4** (8) | **3** (8) |
| *Rpv1-Rpv10* | *Run1-Ren3-Ren9* | Coliris, Opalor, Lilaro, Sirano, Selenor | **2** (2) | **2** (2) | **2** (2) | **2** (2) | **3** (3) | **3** (9) | **1** (8) | **7** (12) |

**Table 2.**
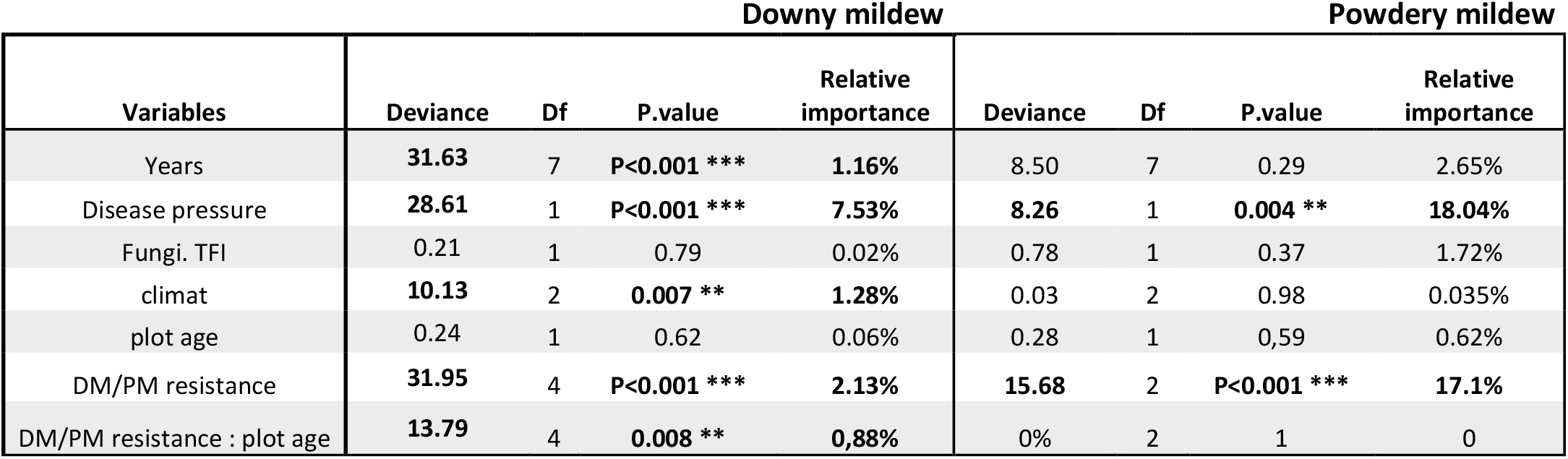
Analysis of deviance based on likelihood ratio tests for the GLM fitted to binary disease severity (high vs low) on leaves. For each term, the number of degree of freedom (Df), the percentage of null deviance explained per degree of freedom (Relative importance estimated as a partial pseudo-R^2^ per Df) and the p-value are reported.

**Table 2.**
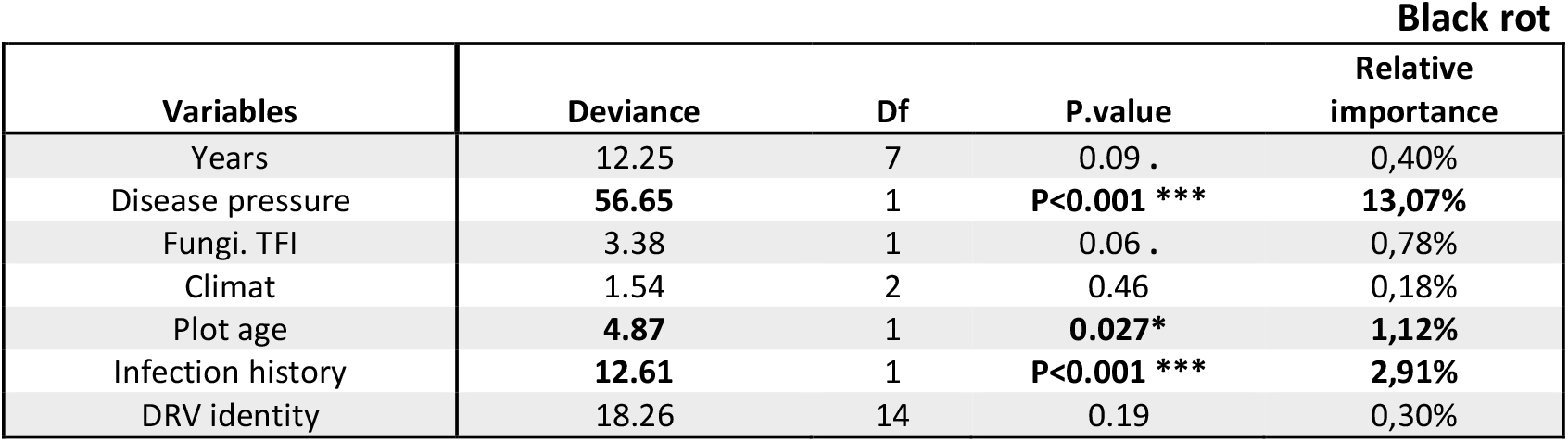
Analysis of deviance based on likelihood ratio tests for the GLM fitted to binary disease severity (high vs low) on leaves. For each term, the number of degrees of freedom (Df), the percentage of null deviance explained per degree of freedom (Relative importance estimated as a partial pseudo-R^2^ per Df) and the p-value are reported.

| Black rot |  |  |  |  |
| --- | --- | --- | --- | --- |
| Variables | Deviance | Df | P.value | Relative importance |
| Years | 12.25 | 7 | 0.09 . | 0,40% |
| Disease pressure | **56.65** | 1 | **P<0.001 \*\*\*** | **13,07%** |
| Fungi. TFI | 3.38 | 1 | 0.06 . | 0,78% |
| Climat | 1.54 | 2 | 0.46 | 0,18% |
| Plot age | **4.87** | 1 | **0.027\*** | **1,12%** |
| Infection history | **12.61** | 1 | **P<0.001 \*\*\*** | **2,91%** |
| DRV identity | 18.26 | 14 | 0.19 | 0,30% |

In France, the cultivation of resistant grapevine varieties has increased steadily over recent years. The total area planted with DRVs nearly tripled within four years, increasing from 1184 ha in 2021 to 3107 ha in 2025. The OSCAR observatory captures part of this expansion, accounting for 6.8% of the total area planted with DRVs in 2021 (81.6), and 4.6% in 2025 (127 ha) (**Figure 1B**).

Although the observatory includes a limited number of plots, varietal diversity was consistently higher than that observed across France. In 2021 and 2025, diversity remained consistently higher in OSCAR (Shannon: 1.69–1.74; Simpson: 0.766–0.801) compared to France (Shannon: 1.39–1.41; Simpson: 0.684–0.694). This result from the dominance at the national scale of two varieties, Souvignier gris (*Rpv3*.*2, Ren3–Ren9*) and Floreal (*Rpv1–Rpv3*.*1, Run1–Ren3–Ren9*), covering respectively 32% and 30% of the DRVs area in 2025. By contrast, they represents 13.9% (Souvignier gris) and 22.5% (Floreal) of the observatory surface. In the opposite, the so-called Bouquet varieties are overrepresented (25% of the total monitored area, representing 98.8% of Bouquet varieties area at the national level) mainly because most of these varieties are not yet officially registered and still at the experimental stage. (**Figure 1)**. In summary, the observatory represents a well-balanced and heterogeneous proportion of resistant grapevine varieties cultivated in France.

### Disease resistant grapevine varieties ensure powdery and downy mildew control with (much) less fungicide use

Data collected over the past 8 years through the OSCAR observatory provide the first insights into the integration of DRVs across French vineyards.

First, the cultivation of DRVs across a wide range of agro-climatic contexts has resulted in an average 79% reduction in the fungicide TFI compared to the national average recorded for susceptible varieties in 2019 (mean in OSCAR = 2.03; Reference Agrest 2019 = 10.1) (**Figure 2A**). While the reduction in fungicide TFI remained highly significant across all years, its magnitude is lower in high-disease-pressure years (*e*.*g*. 3.12 in 2024, corresponding to a 69.1% reduction) than in low-pressure years (*e*.*g*. 0.8 in 2019, corresponding to a 92.1% reduction). Notably, the fungicide TFI in OSCAR remained in 2025 close to the 2024 level, although climatic conditions were less conducive to disease development.

Importantly, this drastic reduction in fungicide use does not result in higher PM and DM severity that remain well controlled on DRVs. Since 2018, the absence of PM symptoms in 98% of plots indicates a near-complete disease control across the observatory. This likely reflects the widespread deployment of the *Run1* factor that confer a total resistance to PM. By contrast, symptoms of DM are more frequently observed, reflecting the quantitative nature of the resistance factors deployed. Nevertheless, its control remains effective: the mean annual probability of observing an high DM severity on leaves (*i*.*e*. score ≥ 2) is 0.15. In 2024, a year highly conducive to DM in France, 80% of the OSCAR plots exhibited less than 5% symptom severity on leaves (**Figure 2B**).

Because most vineyard plots in the observatory were still young, disease symptoms were primarily observed on leaves, as grapevines generally begin producing clusters only after 2–3 years. Among plots with observations available on both leaves and clusters, the probabilities of detecting high disease severity on clusters given its presence on leaves was 0.43 for DM and 0.63 for PM.

### Genetic resistance combination to mildews displays different efficacity and durability

The OSCAR observatory monitored 30 cultivars carrying distinct genetic resistance combinations in 2025. To assess their effectiveness and durability, we fitted binomial GLMs to analyze how the probability of high disease severity depends on local disease pressure, agro- climatic context, TFI, and the interaction between genetic resistance combinations and plot age.

Overall, powdery mildew was rarely detected in the observatory (**Figure 2B**). The resistance factors deployed emerged as the most significant variable affecting probability of high leaf severity (relative importance 17.1%; p<0.001). Three resistance factors are currently deployed in DRVs, namely *Run1, Ren3-Ren9*, or the combination *Run1-Ren3-Ren9*. The probability of observing high PM severity was null in DRVs carrying *Run1*, either alone or in combination with *Ren3*-*Ren9*, confirming the total resistance provided by *Run1*. The few cases of high PM severity were restricted to varieties carrying *Ren3*-*Ren9* only, although no significant differences were detected between these three genetic resistance combinations (**Figure 3B**). Local PM pressure is the second main factors influencing the probability of observing high PM severity on grapevine leaves (relative importance 18.04%; p=0.004). The non-significant interaction between the genetic resistance combinations and plot age suggest that all genetic resistance combinations remain effective over time (**Table 2**).

**Figure 3.**
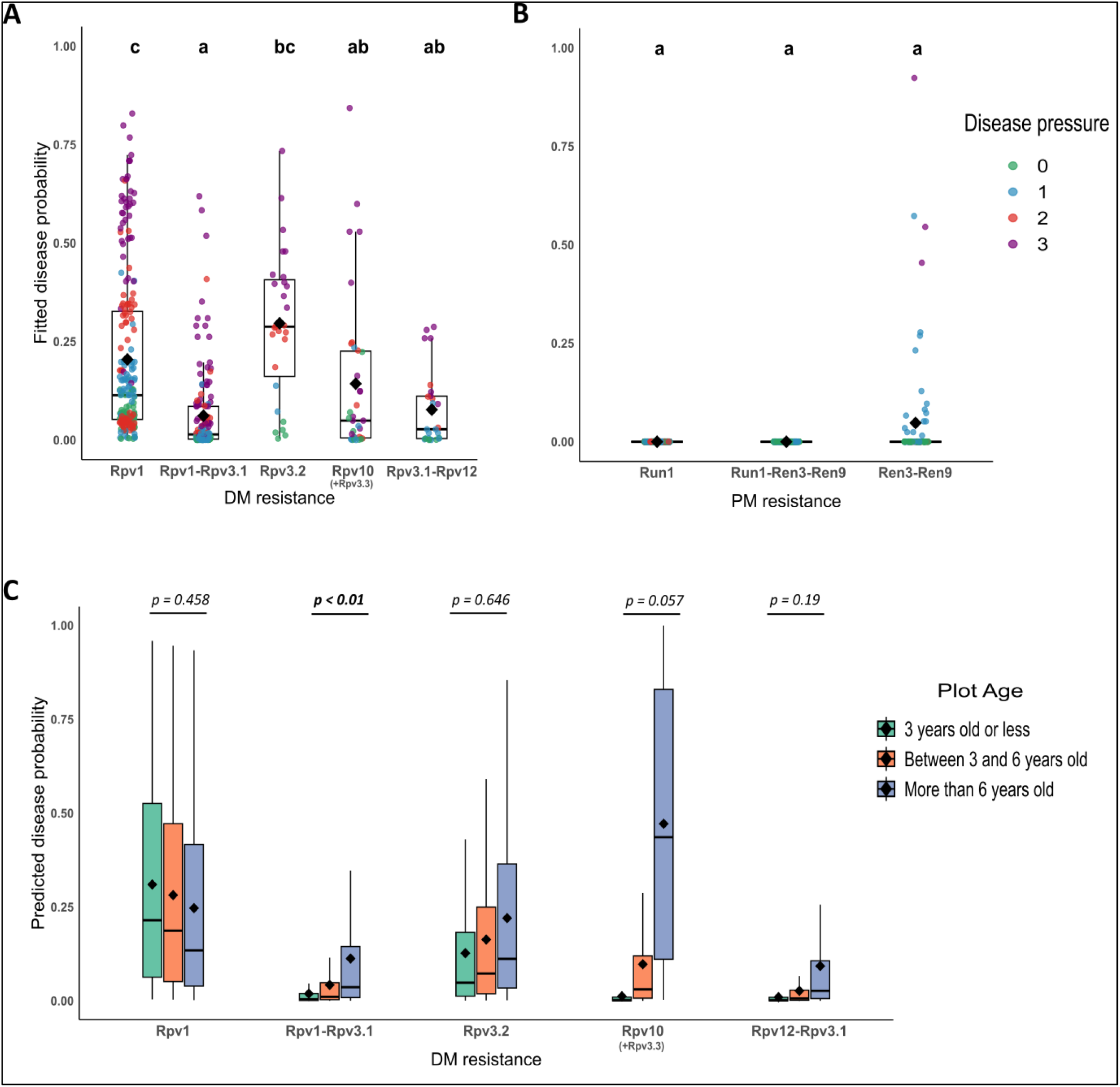
Effects of resistance factors and plot age on the probability of high downy and powdery mildew severity in the OSCAR observatory. **A, B)** Probability of downy mildew (A) and powdery mildew (B) high severity per resistance factor, calculated for each year and climate, at average plot age and mean fungicide dose. Colors indicate pathogen pressure on neighboring susceptible vines (from 0 = no pressure to 3 = high pressure). Different letters indicate significant differences between resistance factors according to Tukey’s HSD post-hoc test (P<0.05). **C)** Predicted downy mildew high severity probability across three plots age classes for each resistance factor; powdery mildew is not shown due to non-significant variation over years.

**Figure 4.**
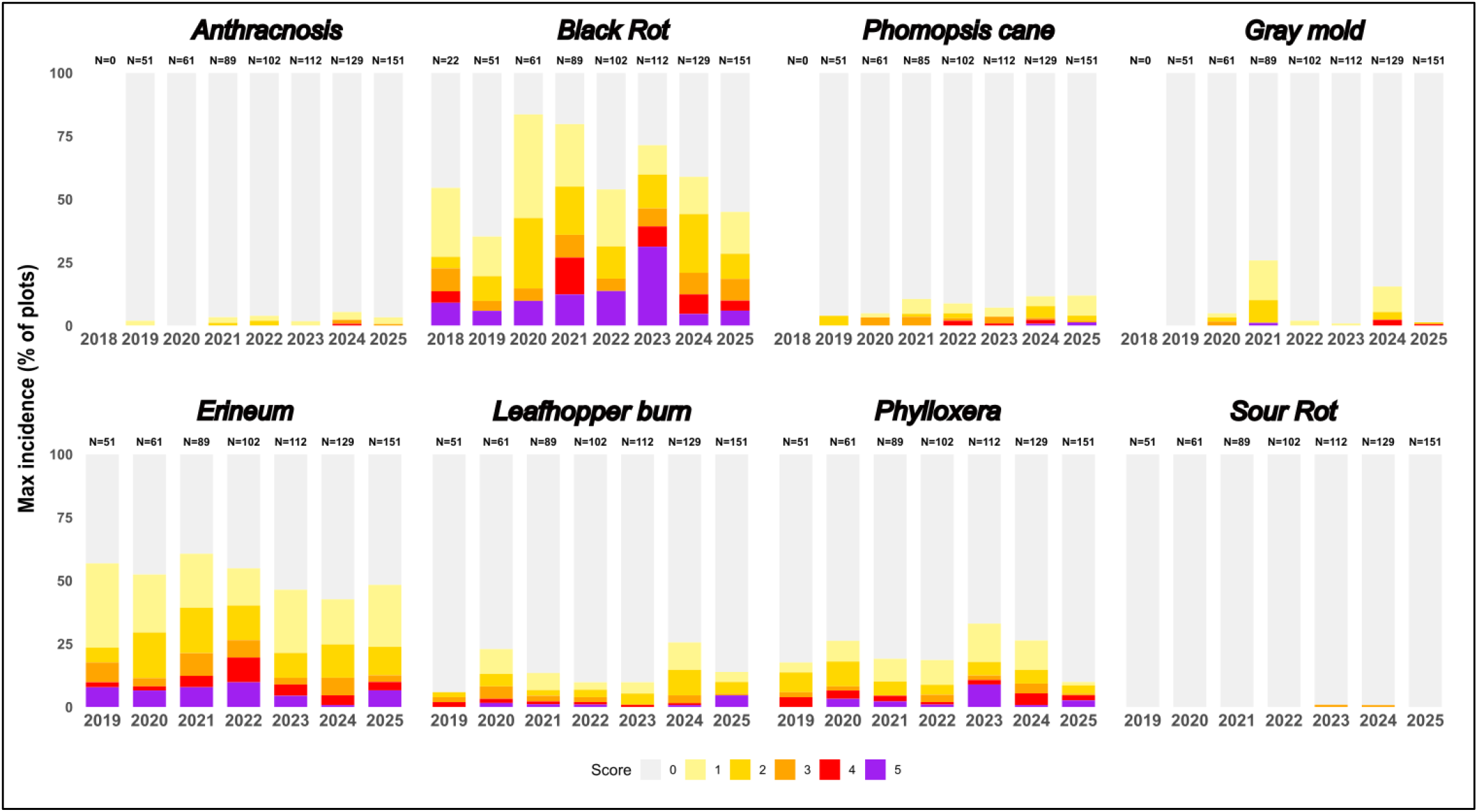
Incidence of the major grapevine pests and diseases monitored in the OSCAR observatory, excluding mildews. Maximum Incidence (Frequency of plant with at least one symptom per plot) of fungal disease (anthracnosis, black rot, phomopsis cane, and grey mold) and insect relative disease (erineum mite, leafhopper damage, foliar phylloxera, and sour rot) on leaves, clusters and/or stems on plots surveyed at veraison from 2018 to 2025. Incidence and severity were calculated for both leaves and clusters, and the maximum value between the two organs is shown. The histograms report the percentage of the plots in each class (from 0 to 5, for score correspondence see methods), the number above represents the total of plots recorded for each year

**Figure 5.**
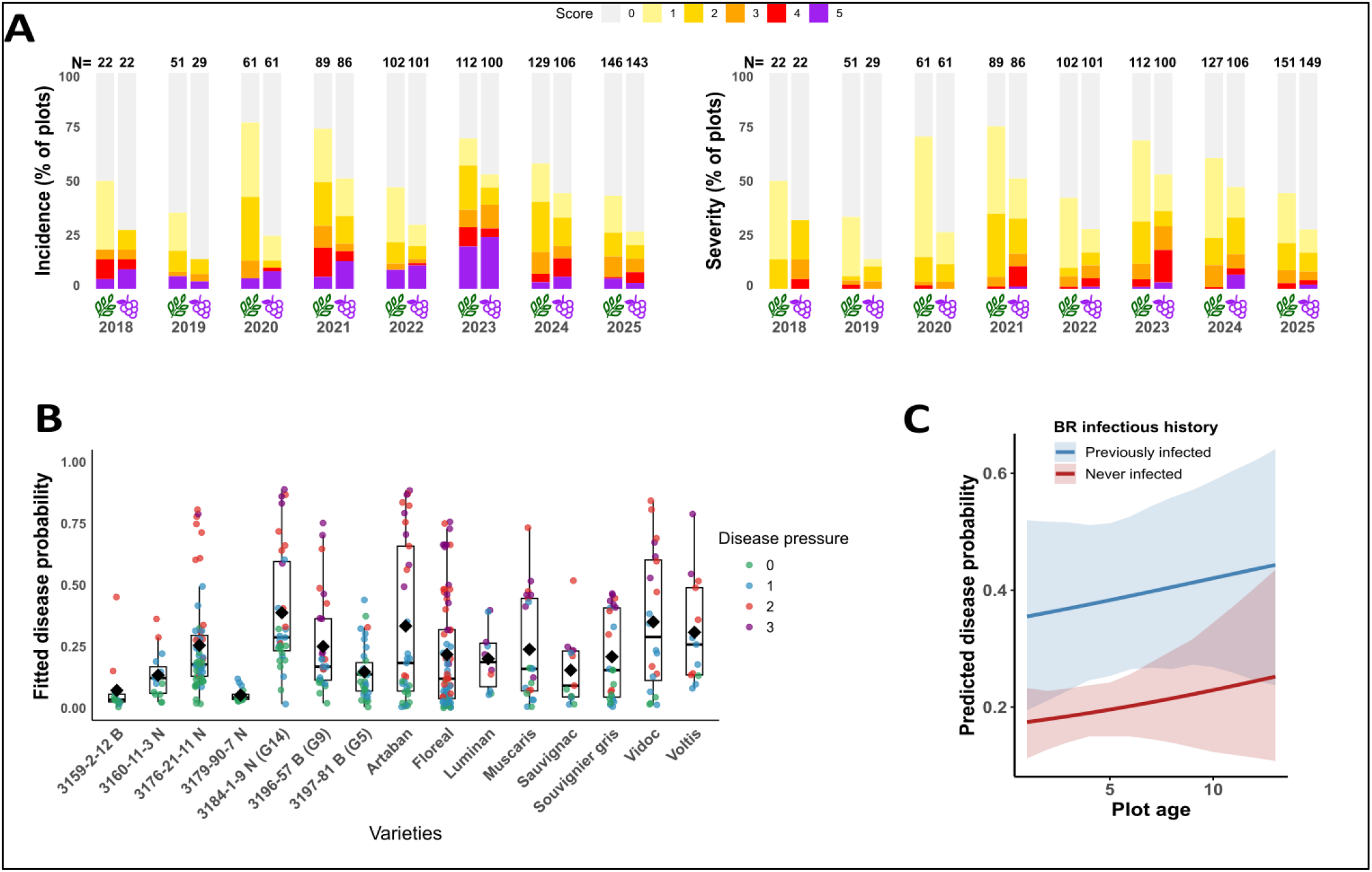
Black rot incidence and severity in the OSCAR observatory. **A)** Incidence (frequency of plant with at least one symptom per plot) and severity (intensity of symptom) of black rot on leaves and clusters in plots surveyed at the veraison or harvest stages since 2018. The histograms report the percentage of the plots in each class (from 0 to 5, for score correspondence see methods), the number above represent the total of plot recorded for each year and each organ. **B)** Probability of high black rot severity for each DRV, calculated for each year and climate, at average plots age and mean fungicide dose. Colors indicate pathogen pressure on neighboring susceptible vines (0 = no pressure to 3 = high pressure). Statistical differences among resistance factors were tested using Tukey HSD test on the model described in methods. **C)** Effect of plot age and prior infection status history on high black rot severity. For each age value and disease-history condition, the 2.5th and 97.5th percentiles of the bootstrap distribution of predicted probabilities were used to combination 95% confidence intervals (see methods).

In contrast to PM, downy mildew was more frequently detected in the observatory. The probability of high DM severity on leaves was primarily driven by local disease pressure, year effect, and the genetic resistance combination deployed. Local DM pressure (relative importance 7.53% ; p<0.001) and year effect (relative importance 1.16%; p < 0.001), consistent with the well-established practitioner knowledge of years more conducive to DM epidemics, were major determinants. The high level of disease pressure significantly increased the marginal probability of high DM severity compared to low pressure (13.5% ± 5% vs. 1.5% ± 0.8%). The genetic combination deployed against downy mildew was also an important driver of high DM severity (relative importance 2.13%; p < 0.001). Among those, *Rpv1* was the least effective, followed by *Rpv3*.*2, Rpv10, Rpv1-Rpv3*.*1*, and *Rpv3*.*1-Rpv12* (**Figure 3A**). Importantly, the probability of high DM severity significantly increased with plot age for the *Rpv1-Rpv3*.*1 g*enetic resistance combination. This effect is nearly significant for the varieties carrying *Rpv10* and *Rpv3*.*1-Rpv12* (**Figure 3C**). These results suggest a decline in the effectiveness of these resistance factors over time. Finally, agro-climatic context also had a significant effect: the deployment of DRVs in the Mediterranean climat significantly reduced the marginal probability of DM high severity compared to the Oceanic one (4.5% [4.1–4.95] vs. 7.8% [7.15–8.07]; p=0.007).

### Dynamics of pests and diseases not targeted by the resistances

Incidence and severity of main grapevine pest and diseases beyond downy and powdery mildew were also monitored within the OSCAR observatory. Four fungal diseases (anthracnosis, black rot, phomopsis cane, and grey mold) and four insect-related diseases (erineum mite, leafhopper damage, foliar phylloxera, and sour rot) were recorded across the monitored plots (**Figure 4**).

Only black rot (BR) shows increasing and concerning levels of incidence and severity. Specifically, BR severity has been rising in OSCAR, with more than 35% of plots showing at least moderate severity >5% on leaves in both 2023 and 2024, and 10% of plots exhibiting at least severe symptoms (>10%) in 2024 on clusters where BR is particularly damaging for the harvest (**Figures 4-5**). Similar to DM, statistical analyses identified local pathogen pressure as one of the main driver of high BR severity on leaf (relative importance 13.07% ; p<0.001). Specifically, high disease pressure significantly increased the marginal probability of high BR severity on leaf compared to low disease pressure (46.7% ± 9.8% vs. 16.8% ± 4.9).

The probability of high BR severity on leaves significantly increased by approximately 0.6 percentage points per year as plot age increased (p=0.027). Prior infection further compounded this risk (p<0.001). For plots without previous infection, the probability of high severity rose from 0.18 in young plots (≤3 years old) to 0.22 in older plots (> 6 years old). For plot having already experienced BR infection, they nearly doubled, reaching 0.35 in young plots and 0.41 in older ones, highlighting the opportunistic nature of the disease. Interestingly, a slight, non-significant reduction in probability of high BR severity was observed with fungicide application, suggesting a biologically consistent trend. Additionally, minor, non-significant, interannual variations were detected, indicating that some years are more conducive to disease development than others. The identity of the DRV had no significant impact on the probability of high BR severity, suggesting that cultivars display comparable susceptibility to BR. Overall, these results indicate that environmental conditions, prior black rot infection history in the plot and plot age are the primary determinants of BR severity, outweighting the effects of fungicide treatment, year, and cultivar (**Figure 5**)

## Discussion

The early establishment of the OSCAR observatory, concurrent with the first deployments of resistant grapevine varieties in France, represents a valuable opportunity to understand the full range of changes induced by resistance deployment, both for disease management and pathogen adaptation. The alert system associated with the observatory has already proven effective: the annual field sampling combined with laboratory phenotyping experiments enabled the several early detections of the emergence of resistance-breaking strains (Dvorak et al., 2025; Paineau et al., 2022; Pelissier et al., 2025). These isolated cases prompted rapid feedbacks and technical supports to the respective winegrowers for DRV management (‘OSCAR observatory, 2026’). Here, we focus instead on the second pillar of the observatory, the long-term monitoring of the DRV in real production situations, which had not been previously analyzed.

Since 2018, the use of grapevine DRVs enables a drastic reduction in fungicide applications (- 80%) in a wide range of production situations, while maintaining effective control of both downy and powdery mildews. Other grapevine diseases and pests remain well controlled on DRVs, with the notable exception of black rot, exhibiting significant increases in both incidence and severity over the years. While grapevine DRVs have previously proven effective in controlling powdery and downy mildews on leaves and clusters in greenhouse and experimental conditions (Calonnec et al., 2013; Miclot et al., 2022; Salotti et al., 2022), this study is the first one evaluating their long-term performance in the field in a wide range of conditions. Beyond viticulture, the OSCAR observatory also represents a unique dataset documenting the efficacy and durability of varietal resistance in perennial crops, addressing a major gap regarding long-term, field-scale evaluations (Van der Graaff, 1983).

For powdery mildew, the observatory includes grapevine DRVs carrying both complete (*Run1*; Feechan et al., 2013) and partial (*Ren3* and *Ren9*; Zendler et al., 2021) resistance factors. The disease was absent in 98% of the 650 observations realized on leaf within the observatory due to the widespread presence of the *Run1* factor, which confers a complete resistance and has, to date, not been overcome in Europe (Sosa-Zuniga et al., 2022). Powdery mildew was exclusively detected on DRVs carrying *Ren3* and *Ren9* which is consistent with the partial nature of these resistance factors (Sosa-Zuniga et al., 2022; Zendler et al., 2021). The durability of the DRVs against powdery mildew over time suggest that no breakdown of *Run1, Ren3*, or *Ren9* occurred yet in Europe. However, a continued vigilance is required as such breakdowns have already been observed in North America, where these resistance factors originate (Agurto et al., 2017; Feechan et al., 2013).

Resistance breakdowns in DRVs have so far been reported in Europe only for downy mildew. However, downy mildew remains well controlled within the OSCAR observatory. Even in 2024, a year highly conducive to disease development, only 7% of monitored plots showed more than 5% of severity on leaves at veraison stage. Nevertheless, signs of loss of efficacity with plot age have been detected for some genetic combination, particularly in ResDur1 variety (*Rpv1–Rpv3*.*1*), and to a lesser extent in varieties carrying *Rpv10* or *Rpv3*.*1-Rpv12*. This suggests the emergence of strains adapted to these genetic combinations. Although rare, *Rpv1*-breaking strains have already been reported so far in France (Pelissier et al., 2025) and in Réunion island (Martinez et al., 2025). However, the efficacy of *Rpv1* alone appears stable over time in the observatory, showing no significant decline with plot age. Similarly, the efficacy of *Rpv3*.*2*, the second most widely deployed resistance factor in France, has remained stable, consistent with the single documented case of its breakdown in Germany to date (Gouveia et al., 2024). In contrast, it is now well established that the large-scale cultivation of hybrid grapevine varieties carrying the *Rpv3*.*1* resistance factor during the 20^th^ century has contributed substantially to the emergence of *Rpv3*.*1* breaking strains across Europe (Delmotte et al., 2014; Di Gaspero et al., 2012). It may explain the reduced efficacy observed in the OSCAR observatory for ResDur1 varieties (*Rpv1–Rpv3*.*1*) and, to a lesser extent, for those carrying *Rpv3*.*1-Rpv12*. The scarcity of varieties carrying *Rpv12* alone in the observatory prevents a direct assessment of its long-term durability. Recent studies describe *Rpv12* as a highly effective, nearly complete resistance (Chitarrini et al., 2020; Dvorak et al., 2025; Paineau et al., 2024; Wingerter et al., 2021). This may explain why *Rpv12* exerts a dominant effect in pyramids like *Rpv12-Rpv3*.*1*, effectively masking the breakdown of *Rpv3*.*1* in most *P. viticola* populations.

While the erosion of resistance efficacy observed in pyramided varieties containing *Rpv3*.*1* was expected (Paineau et al., 2024; Pelissier et al., 2025; Peressotti et al., 2010), the decline in performance of *Rpv10* revealed in this study is more unexpected and deserves further investigation. Indeed, even if monogenic varieties carrying *Rpv10* (e.g. Muscaris) were grouped in our analysis with varieties combining *Rpv10* and *Rpv3*.*3* (e.g. Cabernet Cortis, Monarch, Divico), the observed increase in disease severity within this group cannot be solely attributed to the breakdown of *Rpv3*.*3*, already observed in Europe, as this resistance factor provides only a limited level of protection (Di Gaspero et al., 2012; Foria et al., 2018; Wilkerson et al., 2025). Rather, our observations suggest a potential on-going adaptation of *P. viticola* populations to *Rpv10* in France. Consequently, close monitoring for early signs of *Rpv10* breakdown is highly warranted, especially since such events have already been documented in Germany and Hungary (Heyman et al., 2021; Paineau et al., 2022).

Our result also highlights a notable increase of black rot in DRV plots over the years. This pattern is likely explained by the drastic reduction in fungicide applications in DRVs, made possible by resistance to powdery and downy mildew the two main targets of chemical control in grapevine (Amiot et al., 2024; Szabó et al., 2023). It is well established that treatments against mildews also indirectly suppressed black rot (Szabó et al., 2023). However, our results indicate that additional factors could play a major role. In particular, the infection history of a plot strongly influenced the risk of severe black rot symptoms in subsequent years. These results confirm that black rot is promoted by reduced fungicide protection and the presence of established inoculum in the plot (Hoffman and Wilcox, 2002). Another contributing factor may be the ecological niche left vacant by powdery and downy mildew on grapevine leaves, which could facilitate the emergence of black rot. Examples of fungal competition between pathogens have been documented in the literature, where the suppression of one species (following the deployment of an host resistance) favor another fungal pathogen species (Zhan and McDonald, 2013). The increase of black rot prevalence in the observatory could also reflect indirect consequences of the breeding process. Similar patterns have been described in other crops, such as wheat, where the selection of genomic regions improving yield and rust resistance inadvertently introduced alleles associated with higher susceptibility to *Septoria tritici* blotch (Arraiano and Brown, 2017). Such linkage effects illustrate how breeding for specific traits can unintentionally favor the emergence of secondary pathogens by modifying plant defense pathways or metabolic balances (Brown, 2015). However, given the complex breeding history of resistant grapevine varieties from diverse Asian and American backgrounds (Merdinoglu et al., 2018), and the similar levels of black rot susceptibility observed across varieties (**Figure 5**), this hypothesis is not strongly supported by the available evidence. Future breeding strategies aiming to add black rot resistance to DRV (Bettinelli et al., 2023) should be complemented by agronomic practices reducing inoculum, such as the removal of mummified berries (Hoffman et al., 2004).

Varietal resistance is a cornerstone of sustainable agriculture (Juroszek and von Tiedemann, 2011; Zhan et al., 2014). The high evolutionary potential of plant pathogens makes the long-term monitoring of resistance performance under field conditions essential to ensure their durable management and guide breeding strategies. In annual crops, where crop rotation enables high flexibility, monitoring is often conducted in collaboration with technical institutes, allowing rapid feedbacks to growers and swift adjustments of varietal choices following breakdowns. Examples include sugar beet in France, wheat rusts via RustWatch (“RustWatch,” 2025), and potato late blight through the EuroBlight network (“Euroblight” 2025). In perennial crops, where adaptive management options are inherently limited, large-scale monitoring networks remain surprisingly less developed, despite their instrumental role in guiding resistance breeding strategies, as illustrated by the European Vinquest project on apple scab (Patocchi et al., 2020).

Within this framework, the OSCAR observatory provides a unique plateform to assess the performance of resistant grapevine in production situations over cropping seasons. Given the recent and still limited deployment of resistant grapevine varieties in France, the identification of several resistance breakdowns are particularly concerning (Paineau et al., 2024; Pelissier et al., 2025). These findings emphasize that resistance management must be dynamic, integrating both genetic and agronomic strategies to mitigate risk of breakdown and prolongate the effectiveness of existing resistance factors (Brown, 2015; REX Consortium et al., 2016). A textbook example concerns the *Rpv3*.*1* gene; its widespread deployment has led to the recurrent emergence of adapted *P*.*viticola* strains in several European regions (Delmotte et al., 2014; Di Gaspero et al., 2012; Paineau et al., 2024). Beyond the loss of efficacy itself, *Rpv3*.*1* may also act as a stepping stone for pathogen adaptation in pyramided varieties, thereby compromising their durability (Zaffaroni et al., 2024). As such, its inclusion in future breeding programs should be avoided.

An effective approach to manage the durability of resistance genes is to combine the deployment of resistant grapevine varieties with other agronomic levers, such as varietal mixtures (Mundt, 2002; Zaffaroni et al., 2024), prophylaxis (Poeydebat et al., 2022), or the application of natural or phytosanitary products(Fournier et al., 2025; Zhang et al., 2017). Maintaining a limited fungicide protection on DRVs can indeed slow down pathogen adaptation by limiting population sizes (Zaffaroni et al., 2025), especially during years of high disease pressure (McDonald and Linde, 2002). Noteworthy, the OSCAR observatory recently recommended to maintain a baseline phytosanitary protection on DRVs that experienced breakdowns in 2024 in southeastern France (Pelissier et al. 2025), even during years with low epidemic pressure (‘OSCAR observatory, 2026’). This proactive recommendation likely explains the small TFI difference observed between 2024 (high DM pressure) and 2025 (lower DM pressure). Finally, the increasing prevalence of black rot in the OSCAR observatory further supports this recommendation.

Overall, the results emerging from the OSCAR observatory highlight the importance of long-term, coordinated monitoring networks to track pathogen adaptation in response to the deployment of resistant cultivars. Although such observatories are challenging to design and maintain (Soubeyrand et al., 2024), they are necessary for proactively managing resistances. Today, the OSCAR network is expanding to the main European wine-producing countries through an international coordination effort, thereby extending surveillance and strengthening early detection of grapevine resistance breakdowns. Such large-scale initiatives are crucial to guide future breeding programs and maximize the prospects of sustainable disease management in perennial crops.

## Acknowledgements

The authors would like to thank the staff of their research unit (SAVE) for their assistance in manuscript preparation and for valuable discussions. We are also grateful to all the partners of the OSCAR network for ensuring the annual monitoring of the plots. Specifically, we would like to acknowledge BNIC (Bureau National Interprofessionnel du Cognac), CIVA (Conseil Interprofessionnel des Vins d’Alsace), CIVC (Comité Interprofessionnel des Vins de Champagne), and CIVL (Conseil Interprofessionnel des Vins du Languedoc), as well as the Chambres d’agriculture of Allier, Aude, Bouches-du-Rhône, Centre-Val de Loire, Dordogne, Drôme, Gard, Gironde, Hérault, Pays de la Loire, Pyrénées-Orientales, Vaucluse, and Vienne. We are also gratefull to the cooperative wineries Alliance Aquitaine, Cave du Marmandais, Terres de Valdèze, Vignerons Ardéchois, Vignerons de Buzet, Vignerons de Landerrouat, Vignerons de Tutiac, and Vinovalie. We also thank IFV (Institut Français de la Vigne et du Vin), Agrobiopérigord, SRDV (Société de Recherche et de Développement Viticole), and ICV (Institut Coopératif du Vin), along with the INRAE experimental viticulture units of Bordeaux, Colmar, Pech Rouge, and Montpellier, the Lycée viticole de Rodilhan, and all participating winegrowers involved in the project.

## Author Contribution

RP, ASM, LD, FD, and FF contributed to the conceptualization. RP wrote the original draft. RP, ASM, LD, FD, and FF contributed to the review and editing. RP, LM, and IDM contributed to data curation and formal analysis. ASM, LD, FD, and FF contributed to funding acquisition and project administration. RP and FF contributed to the methodology.

